# Cyclin B3 is a dominant fast-acting cyclin that drives rapid early embryonic mitoses

**DOI:** 10.1101/2023.08.11.553011

**Authors:** Pablo Lara-Gonzalez, Smriti Variyar, Jacqueline Budrewicz, Aleesa Schlientz, Neha Varshney, Andrew Bellaart, Shabnam Moghareh, Anh Cao Ngoc Nguyen, Karen Oegema, Arshad Desai

## Abstract

In many species, early embryonic mitoses proceed at a very rapid pace, but how this pace is achieved is not understood. Here we show that in the early *C. elegans* embryo, cyclin B3 is the dominant driver of rapid embryonic mitoses. Metazoans typically have three cyclin B isoforms that associate with and activate Cdk1 kinase to orchestrate mitotic events: the related cyclins B1 and B2 and the more divergent cyclin B3. We show that whereas embryos expressing cyclins B1 and B2 support slow mitosis (NEBD to Anaphase ∼ 600s), the presence of cyclin B3 dominantly drives the ∼3-fold faster mitosis observed in wildtype embryos. CYB-1/2-driven mitosis is longer than CYB-3-driven mitosis primarily because the progression of mitotic events itself is slower, rather than delayed anaphase onset due to activation of the spindle checkpoint or inhibitory phosphorylation of the anaphase activator CDC-20. Addition of cyclin B1 to cyclin B3-only mitosis introduces an ∼60s delay between the completion of chromosome alignment and anaphase onset, which likely ensures segregation fidelity; this delay is mediated by inhibitory phosphorylation on CDC-20. Thus, the dominance of cyclin B3 in driving mitotic events, coupled to introduction of a short cyclin B1-dependent delay in anaphase onset, sets the rapid pace and ensures fidelity of mitoses in the early *C. elegans* embryo.

## INTRODUCTION

During the division of cultured human somatic cells, the interval from nuclear envelope breakdown (NEBD) to sister chromatid separation/anaphase onset is ∼30-60 minutes in the basal state when there are no stressors that prolong mitosis (Araujo et al., 2016). In these cells, the spindle checkpoint is activated during each cell cycle and contributes to setting basal state mitotic timing (Meraldi et al., 2004). In contrast, during the rapid early embryonic divisions of species such as *Drosophila* and *C. elegans*, the basal NEBD to anaphase onset interval is ∼3-5 minutes and the spindle checkpoint is only activated to delay mitotic progression when problems in chromosome-spindle attachment arise (Essex et al., 2009; Yuan and O’Farrell, 2015). How the accelerated mitotic pace characteristic of these early embryonic divisions is achieved is not clear.

Cyclin B-Cdk1 complexes orchestrate the execution of mitotic events (Crncec and Hochegger, 2019; Lindqvist et al., 2009). Following DNA replication, Cyclin B gradually accumulates to drive a bi-stable Cdk1 activity switch (Hegarat et al., 2016; Hutter et al., 2017). Once active, Cyclin B-Cdk1 complexes phosphorylate cellular substrates to direct chromosome condensation, NEBD, spindle assembly, and chromosome segregation (Crncec and Hochegger, 2019; Lindqvist et al., 2009). In late mitosis, the Anaphase-Promoting Complex/Cyclosome (APC/C), an E3 ubiquitin ligase, targets cyclin B for degradation (Alfieri et al., 2017; Watson et al., 2019), leading to sister chromatid separation/anaphase onset and events associated with mitotic exit (Vagnarelli, 2021). In most metazoans, there are three cyclin B isoforms that activate Cdk1 *in vitro*: cyclins B1, B2, and B3 (Lozano et al., 2012). Of these, cyclin B1 and cyclin B2 are closely related in sequence and in evolutionary origin, whereas cyclin B3 is divergent and shares sequence similarity to A-type cyclins (Gallant and Nigg, 1994; Lozano et al., 2012). Phylogenetic analysis has indicated that cyclin B3 emerged coincident with multi-cellularity (Lozano et al., 2012). Genetic studies have implicated cyclin B3 in the germline and in early embryos (Deyter et al., 2010; Jacobs et al., 1998; Nguyen et al., 2002; Yuan and O’Farrell, 2015). In vertebrates, cyclin B3 expression is largely germline-specific and it has been primarily implicated in female meiosis, where it is essential for the transition from metaphase to anaphase of oocyte meiosis I (Bouftas et al., 2022; Karasu et al., 2019; Li et al., 2019; Zhang et al., 2015).

Here, we analyze the contributions of different cyclin B-Cdk1 complexes to the rapid mitosis of the early *C. elegans* embryo, where NEBD to anaphase onset takes only ∼200s. We show that, whereas embryos with cyclins B1 and B2 support slow mitosis (NEBD to Anaphase ∼ 600s), the presence of cyclin B3 leads to a dominant ∼3-fold acceleration in the pace of mitosis. Cyclin B1/B2-driven mitosis is slower than cyclin B3-driven mitosis largely because the progression of mitotic events is slower. In the context of rapid cyclin B3-driven mitoses, the additional presence of cyclin B1 introduces an ∼1 minute delay, mediated by CDC-20 phosphorylation, in APC/C activation and anaphase onset. Thus, in the early *C. elegans* embryo, the rapid pace of embryonic mitoses is achieved through the dominance of cyclin B3 in driving mitotic events, coupled to introduction of a short cyclin B1-dependent delay that ensure fidelity of chromosome segregation by preventing premature anaphase onset.

## RESULTS AND DISCUSSION

### Cyclin B1 and cyclin B3 independently support CDK-1 activation during early embryonic divisions

To delineate the contributions of cyclin B isoforms (**Fig. 1A; S1A**) to early embryonic divisions in *C. elegans*, we employed RNAi to compare the effects of their inhibition to knockdown of CDK-1. CDK-1 depletion prevented embryos from undergoing mitotic divisions and led to arrest at the one-cell stage with a large polyploid nucleus that arises due to continued DNA replication without intervening mitoses (Edgar and Orr-Weaver, 2001). CYB-3 can be specifically targeted by RNAi, but high sequence identity between CYB-1 and the two CYB-2 isoforms (CYB-2.1 and CYB-2.2 encoded by distinct genes) leads to their simultaneous targeting (referred to as *cyb-1&2(RNAi)*). Whereas simultaneous knockdown of all three cyclins (*cyb-1&2 + cyb-3(RNAi); No CYBs*) lead to an arrest similar to CDK-1 knockdown, expression of either CYB-1&2 or CYB-3 supported mitotic entry (**Fig. 1A-C**; **Fig. S1B**). To evaluate the independent contributions of CYB-1 and CYB-2, we introduced a single copy transgene expressing CYB-1, whose nucleotide sequence was altered to preserve coding information but confer resistance to the dsRNA that targets endogenous CYB-1 and CYB-2, into a *cyb-1*Δ mutant background (**Fig. 1D; S1B-D**); the ability to selectively knockdown CYB-1 versus CYB-2 in the *CYB-RECODE* strain was confirmed by immunoblotting (**Fig. 1E**). Using this approach, we found that CYB-1, but not CYB-2, promoted entry into mitosis (**Fig. 1F**). In contrast to the importance of CYB-1 but not CYB-2 during embryonic mitoses, CYB-1 and CYB-2 functioned redundantly during the meiotic divisions that precede the first embryonic mitosis (**Fig. S2A&B**; (van der Voet et al., 2009)). These results indicate that CYB-1 and CYB-3 are independently capable of supporting CDK-1 activation during early embryonic divisions.

**Figure 1.**
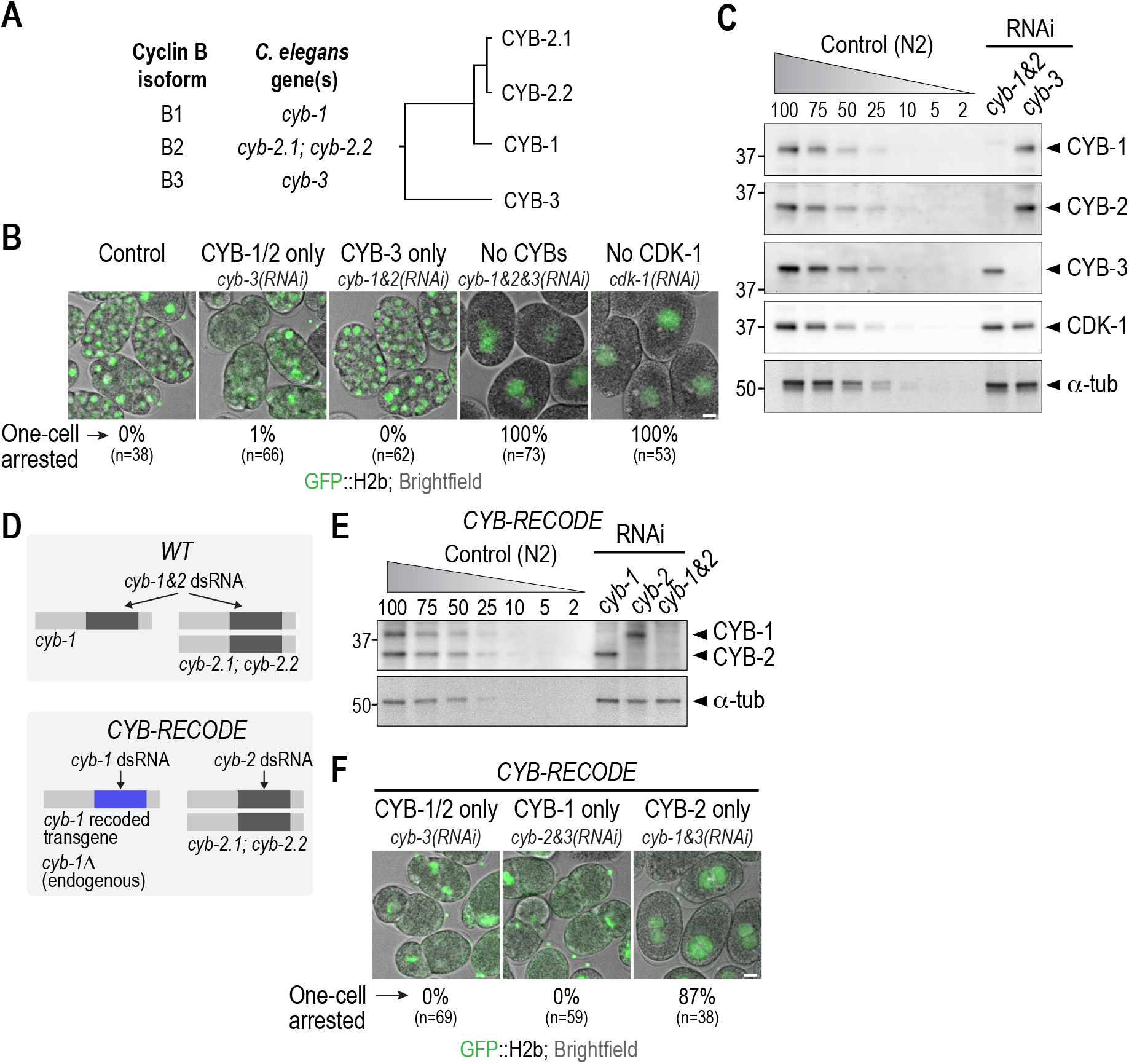
Cyclins B1 and B3 redundantly activate Cdk1 to promote embryonic divisions. **(A)** Phylogenetic tree showing gene names and relationship between *C. elegans* cyclin B isoforms. **(B)** Example images of embryos depleted for the specified proteins. To aid interpretation, the cyclin B(s) present are labeled on top and the RNAi condition employed to generate the specific cyclin B state on the bottom. Quantification of the percentage of interphase-arrested embryos with a single large nucleus is noted below each panel. **(C)** Immunoblot showing depletion of CYB-1&2 or CYB-3 by RNAi. A dilution curve of control (N2) worm extract was loaded to assess depletion efficiency; numbers above lanes indicate percent loading relative to RNAi samples. α-tubulin serves as a loading control. **(D)** Schematic illustrating the principle behind the *CYB-RECODE* system for specific depletion of CYB-1 or CYB-2. **(E)** Validation of the *CYB-RE-CODE* system by immunoblotting, conducted as in panel *(C)*. α-tubulin serves as a loading control. **(F)** Example images of *CYB-RECODE* embryos depleted for the specified proteins, as well as quantification of the percentage of interphase arrested embryos. Scale bars in *(B)* & *(F)*, 10 µm; *n* is the total number of embryos scored per condition.

### Cyclin B1/B2 support slow mitosis, whereas cyclin B3 dominantly drives fast mitosis

We next investigated the effect on mitosis of one-cell embryos of cyclin B isoform depletions. To complement prior surveys conducted using differential interference contrast (DIC) microscopy (Michael, 2016; van der Voet et al., 2009), we compared embryos expressing only CYB-1/2 to embryos expressing only CYB-3 in a strain with fluorescently labeled microtubules and chromosomes. This analysis revealed a striking difference: embryos in which mitosis was driven by CYB-1/2 (*cyb-3(RNAi)*, hereafter referred to as “CYB-1/2-only”) took ∼3 times longer to progress through mitosis than embryos in which mitosis was driven by CYB-3 (*cyb-1/2(RNAi)*, hereafter referred to as “CYB-3-only”; **Fig. 2A,B**). To analyze mitotic progression, we quantified chromosome alignment by measuring the width along the spindle axis of the minimal bounding box that contains all of the chromosomes (termed Chromosome Span, (Cheerambathur et al., 2017)). This analysis revealed that chromosome alignment occurred at a similar rate in control and CYB-3-only embryos but was substantially slower in CYB-1/2-only embryos (**Fig. 2C&D**). Thus, the overall pace of mitotic events including chromosome alignment and anaphase onset is substantially slower in CYB-1/2-only embryos compared to control and CYB-3-only embryos. When CYB-3 is present it overrides the rate set by CYB-1/2 to dominantly drive an ∼3-fold faster mitotic pace.

**Figure 2.**
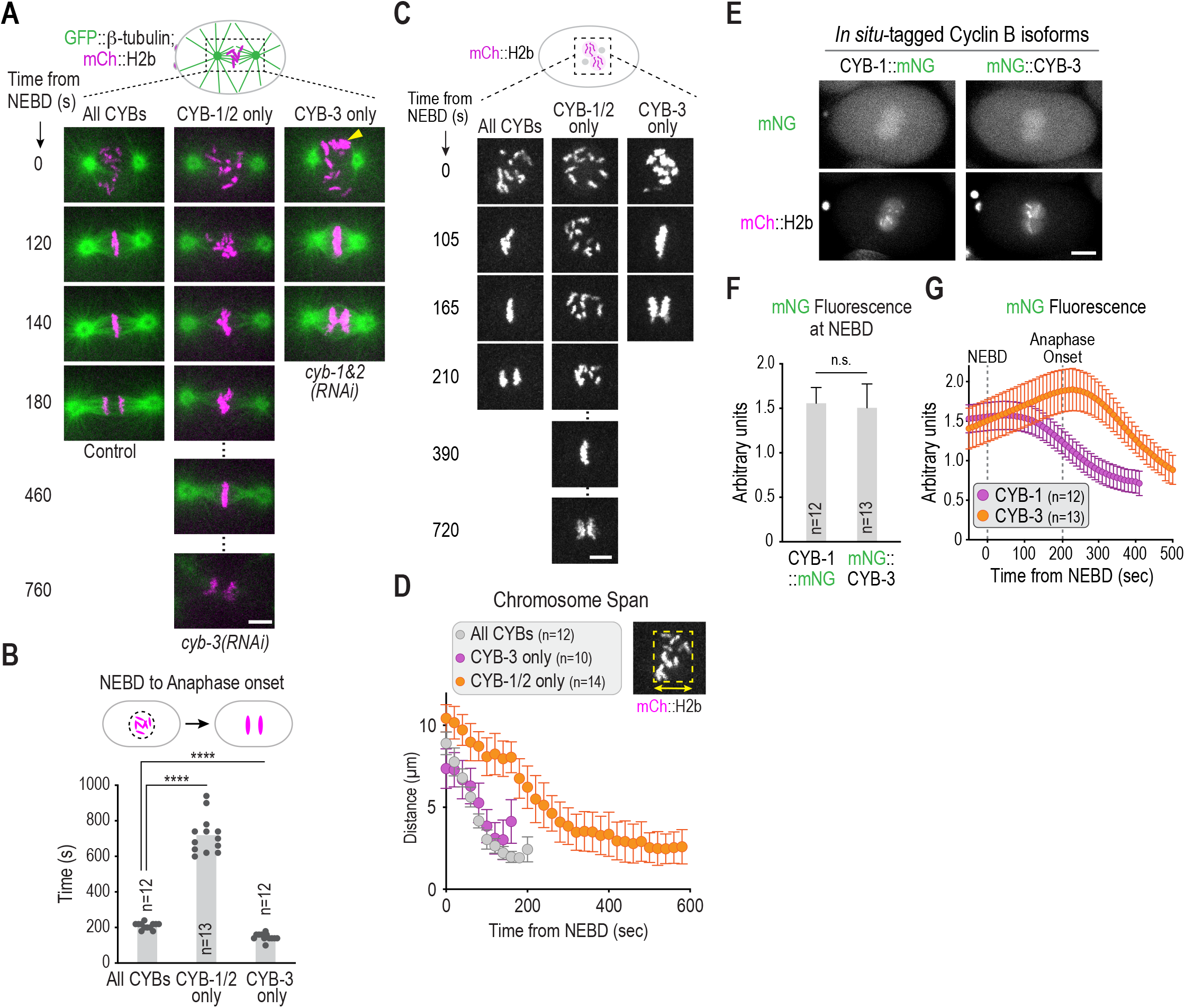
CYB-1/2 supports slow mitosis, whereas CYB-3 dominantly drives rapid mitosis. **(A)** Stills from timelapse sequences of embryos expressing GFP-tagged β-tubulin and mCherry-tagged histone H2b for the specified conditions. To aid interpretation, the cyclin Bs present and supporting mitosis are labeled on top and the RNAi condition employed to generate the specific cyclin B state on the bottom; only the top labels are shown in subsequent figure panels. Yellow arrowhead indicates pronucleus with excess chromatin (see *Fig. S2*). **(B)** Quantification of the nuclear envelope breakdown (NEBD) to anaphase interval for the specified conditions. **(C)** Stills from timelapse sequences of embryos expressing mCherry-tagged Histone H2b for the indicated conditions. **(D)** Quantification of chromosome alignment kinetics by measurement of chromosome span for the indicated conditions. **(E)** Example images of embryos expressing endogenously-tagged cyclin B isoforms. **(F)** Quantification of CYB-1::mNG and mNG::CYB-3 total embryo fluorescence intensity at NEBD. **(G)** Quantification of embryo mNG fluorescence over time for the specified strains expressing *in situ*-tagged cyclin B isoforms (see also *Fig. S2C&D*). In all graphs, *n* is the number of embryos analyzed and error bars are 95% confidence intervals. **** represents p<0.0001 from Mann-Whitney tests; non-significant (n.s.) is p>0.05. Scale bars, 5 µm for panels A & C; 10 µm for panel E.

One explanation for the dominance of CYB-3 is that it represents the predominant cyclin B isoform during early embryonic divisions. To address this possibility, we *in situ*-tagged CYB-1 and CYB-3 at their endogenous loci with mNeongreen (mNG) (Lara-Gonzalez et al., 2019) and quantified the total embryo mNG fluorescence at NEBD (wildtype embryos lacking any mNG fusion were imaged under identical conditions to measure and subtract autofluorescence). This analysis revealed equivalent levels of CYB-1 and CYB-3 in the early embryo (**Fig. 2E&F; S2C&D**), arguing against the dominance of CYB-3 arising from higher abundance relative to CYB-1. Time-lapse imaging of the mNG fusions additionally revealed that CYB-3 is degraded after CYB-1 (**Fig. 2G; S2C&D**), in agreement with prior analysis of cyclin B3 in other systems (Nguyen et al., 2002; Sigrist et al., 1995).

### CYB-3 accelerates mitotic events and APC/C activation without making a significant contribution to reversing inhibitory phosphorylation of CDC-20

The results above indicate that in the early *C. elegans* embryo cyclin B3-driven mitosis is about 3 times faster than cyclin B1/2-driven mitosis. To understand the basis for this difference, we first tested if the slower mitosis in cyclin B1/2-only embryos resulted from activation of the spindle checkpoint, which surveils chromosome-spindle attachments to control anaphase onset (Lara-Gonzalez et al., 2021; McAinsh and Kops, 2023; Musacchio, 2015). Monitoring spindle assembly in embryos expressing GFP::β-tubulin and mCh::H2b revealed that CYB-1/2-only embryos had weaker microtubule density compared to control and CYB-3-only embryos (**Fig. 3A**). In addition, CYB-3-only embryos had shorter spindles, whereas CYB-1/2-only embryos had longer spindles, compared to controls (**Fig. S2E**; (Deyter et al., 2010)). While the spindle checkpoint does not control basal mitotic timing in early *C. elegans* embryos (Houston et al., 2023; Kim et al., 2017), unattached kinetochores resulting from problems in spindle assembly activate the checkpoint to generate checkpoint complexes that extend mitosis by inhibiting APC/C activation (Essex et al., 2009). To determine if checkpoint activation prolongs mitosis in CYB-1/2 only embryos, we co-depleted the checkpoint effector MAD-3 (Essex et al., 2009; Houston et al., 2023; Kim et al., 2017; Lara-Gonzalez et al., 2019). While MAD-3 depletion led to a reduction in the NEBD-anaphase onset interval in CYB-1/2-only embryos, it did not accelerate the rate of chromosome alignment (**Fig. 3B&C**), and the overall mitotic duration remained ∼2.5-fold longer than in controls (**Fig. 3D**). Thus, although the spindle checkpoint is mildly activated in CYB-1/2-only embryos, it is not the primary reason why mitosis is slower compared to controls. In agreement with these findings, evaluation of APC/C activity by measuring the degradation of CYB-1::mNG in living embryos confirmed that APC/C activation is significantly delayed in CYB-1/2-only embryos compared to controls and that APC/C activation is only modestly accelerated by co-inhibition of the spindle checkpoint (**Fig. 3E&F**).

**Figure 3.**
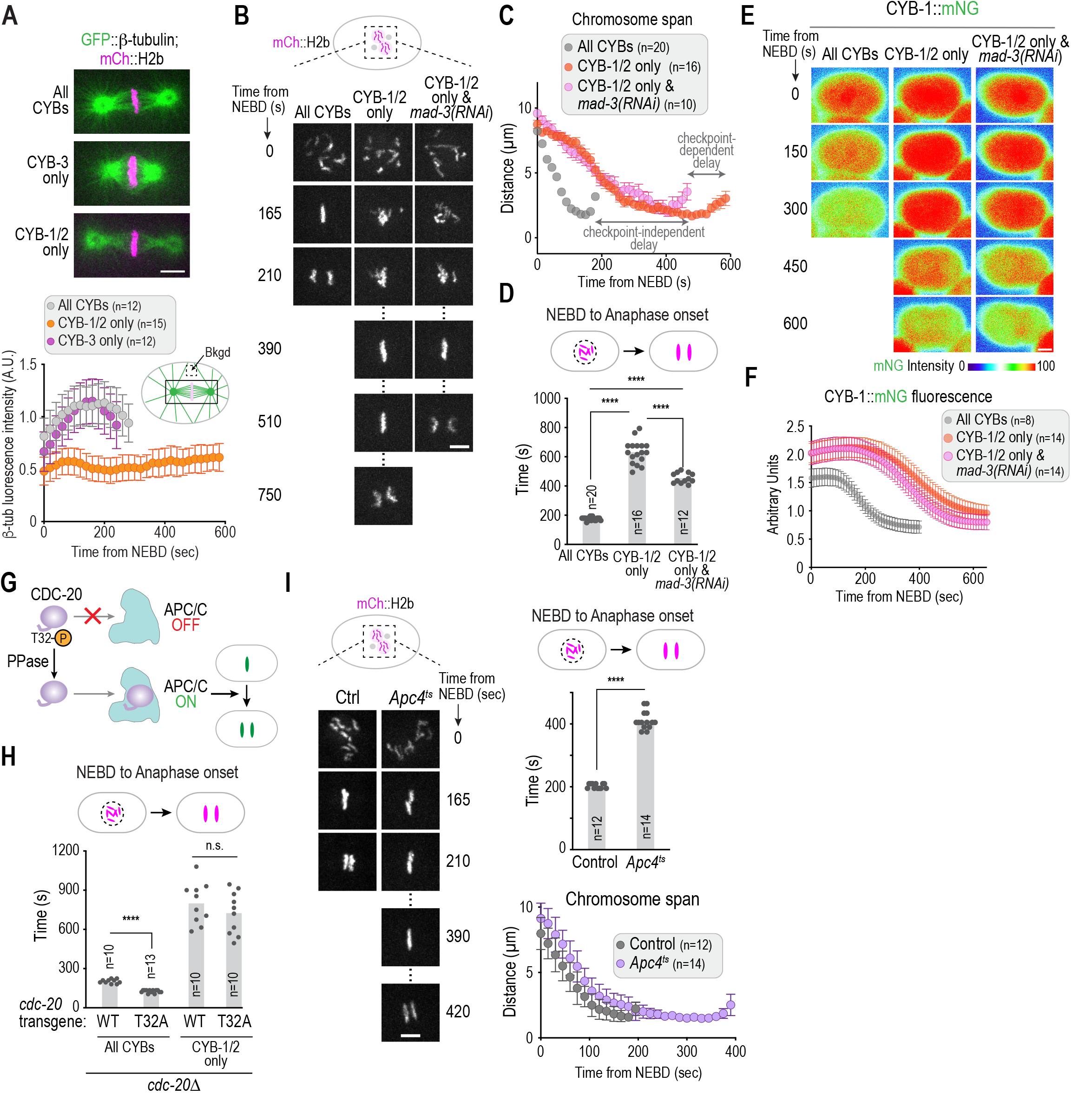
Slow progression of CYB-1/2-driven embryonic mitosis is largely independent of the spindle checkpoint or CDC-20 phosphorylation. **(A)** *(top)* Example images of spindles and chromosomes for the indicated conditions. *(bottom)* Quantification of spindle β-tubulin intensity over time; background fluorescence in a region adjacent to the spindle was subtracted, as indicated by the schematic. **(B)** Stills from timelapse sequences of embryos expressing mCherry-tagged Histone H2b, for the specified conditions. **(C)** Quantification of the chromosome span (*see Fig. 2C&D*) for the specified conditions. **(D)** Quantification of the NEBD to Anaphase interval for the specified conditions. (**E)** Stills from timelapse sequences of embryos expressing mNeogreen-tagged CYB-1. The fluorescence intensity was pseudocolored to aid visualization of change in fluorescence (see also *Fig. S2C&D*). **(F)** Quantification of CYB-1 fluorescence for the specified conditions. **(G)** Schematic describing how phosphorylation of CDC-20 at the Cdk-targetd Thr32 residue controls APC/C activation and anaphase onset. **(H)** Quantification of the NEBD to Anaphase interval in embryos expressing either wild-type or T32A versions of *cdc-20* in a *cdc-20*Δ background. The “*All CYBs”* conditions are the same as for Fig.5B (I) *(left)* Stills from timelapse sequences for wild-type or *Apc4^ts^* (*emb-30(tn377)*) mutant embryos expressing mCherry-tagged Histone H2b. *(top right)* Quantification of the NEBD to Anaphase time interval. *(bottom right)* Quantification of chromosome span over time for the specified conditions. In all graphs, *n* is the number of embryos analyzed and error bars, if shown, are the 95% confidence interval. **** represents p<0.0001 from Mann-Whitney tests; non-significant (n.s.) is p>0.05. Scale bars, 5 µm for panels A, B & I; 10 µm for panel E.

In addition to the spindle checkpoint, APC/C activation can be suppressed by inhibitory phosphorylation of its activating Cdc20 subunit. Biochemical work established that phosphorylation of the N-terminus of Cdc20 family members by Cdks significantly reduces their affinity for the APC/C (Hein and Nilsson, 2016; Labit et al., 2012; Yudkovsky et al., 2000) and structural studies have provided an explanation for the affinity reduction, as the key target sites are in the vicinity of the C-box, the major APC/C binding motif of Cdc20 (Zhang et al., 2016). Prior work in the *C. elegans* embryo has shown that Cdk phosphorylation of a single conserved Cdk-targeted Threonine residue (Thr 32) in the N-terminus of CDC-20 is important for proper mitotic timing (**Fig. 3G**); mutation of this residue to alanine significantly accelerates anaphase onset, whereas mutation to a slower-dephosphorylated serine significantly delays it (Houston et al., 2023; Kim et al., 2017). We therefore considered the possibility that CYB-3 accelerates mitotic progression by reducing the inhibitory phosphorylation of CDC-20 on its T32 residue. However, the NEBD-to-anaphase interval was nearly identical in CYB-1/2 only-embryos expressing wild-type or non-phosphorylatable T32A CDC-20 (**Fig. 3H**), indicating that increased inhibitory phosphorylation of CDC-20 is not a contributor to the delayed APC/C activation in CYB-1/2-only embryos.

Next, we determined whether the reduced rate of chromosome alignment observed in CYB-1/2 only embryos could be a consequence of delayed APC/C activation. For this purpose, we inhibited the APC/C directly using a temperature-sensitive allele of Apc4/EMB-30 (Furuta et al., 2000). At the semi-permissive temperature of 20°C, *Apc4^ts^* caused a 2-fold increase in mitotic duration (**Fig. 3I**), which is close to the ∼2.5-fold increase observed in CYB-1/2-only embryos in which the checkpoint had been inactivated by MAD-3 depletion. However, unlike in the MAD-3 depleted CYB-1/2-only embryos, chromosome alignment occurred with kinetics identical to controls in Apc4^ts^ embryos (**Fig. 3I**).

Collectively, these results indicate that the primary contribution of CYB-3 is to dominantly accelerate mitotic events and the kinetics of APC/C activation. In addition, CYB-3’s contribution to accelerating APC/C activation does not involve reversal of the inhibitory Cdk phosphorylation on CDC-20 and only mildly depends on the spindle assembly checkpoint. These results are most consistent with the notion of CYB-3 being a “fact-acting” cyclin that drives rapid phosphorylation of Cdk1 substrates, when compared to CYB-1/2, thereby accelerating the overall pace of mitosis. In addition to substrates promoting spindle assembly and chromosome alignment, a major Cdk1 substrate is the APC/C, whose phosphorylation releases an autoinhibitory loop, thereby facilitating binding to CDC-20 and substrate targeting (Fujimitsu et al., 2016; Garrido et al., 2020; Qiao et al., 2016; Zhang et al., 2016). While CYB-3 and CYB-1/2 are independently capable of supporting APC/C activation, CYB-3 is able to do it significantly faster and thereby set the duration of mitosis in rapidly dividing embryonic cells.

### CDK-1 association is required for Cyclin B3 to promote rapid embryonic mitosis

The above results suggest that CYB-3 sets the pace of mitosis in early embryos, likely by associating with CDK-1 to form active kinase complexes. To confirm the significance of CDK-1 association, we generated a mutant form of CYB-3 that does not interact with CDK-1. This mutant was designed based on an Alphafold Multimer model of CYB-3 bound to CDK-1 (**Fig. 4A**), which revealed an interaction surface similar to that of cyclin B1 and B2 (Brown et al., 2015), and on prior work that defined a conserved N-terminal helix at the interface in *Xenopus* cyclin B2 that mediates interaction with Cdk1 (Goda et al., 2001). We therefore mutated the key residues in this helix, which correspond to Y112, I116 and Y119 in CYB-3 (referred to herein as CDK-1 binding mutant or CB^Mut^; **Fig. 4A**). We generated strains with integrated transgenes expressing mNeonGreen-tagged versions of WT or CB^Mut^ CYB-3. Immunoprecipitation analysis showed that, while wild-type mNG::CYB-3 co-purified CDK-1, CB^Mut^ mNG::CYB-3 did not (**Fig. 5B**). Similar results were obtained when the CYB-3 – CDK-1 interaction was monitored by co-expression in human cells (**Fig. S3A**).

**Figure 4.**
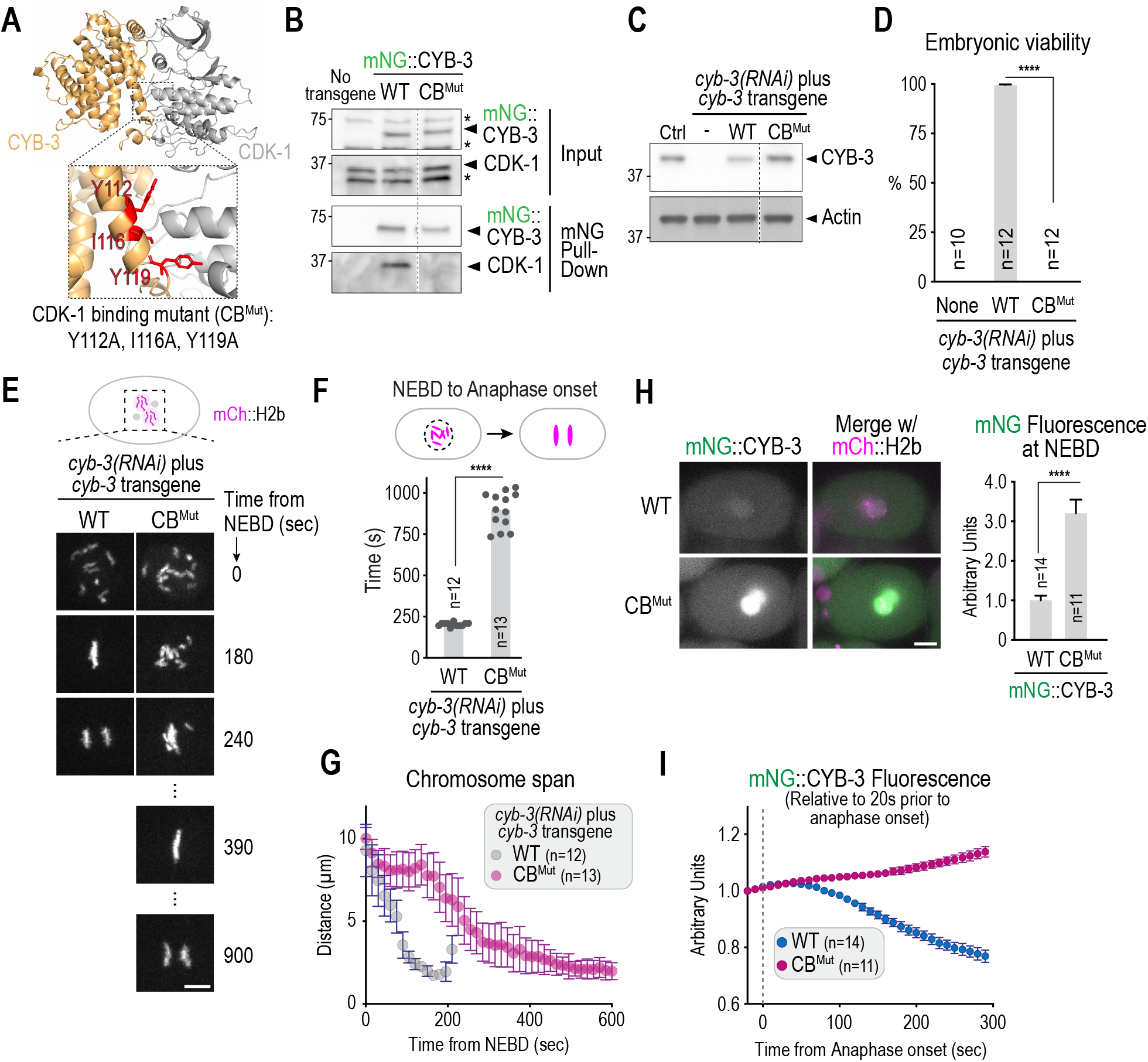
Cyclin B3 accelerates mitotic events by activating CDK-1. **(A)** AlphaFold Multimer model of the *C. elegans* CYB-3 - CDK-1 complex. Mutated interface residues are shown in red. **(B)** Immunoblot showing immunoprecipitation of mNeonGreen-tagged CYB-3 (WT or CB^Mut^) from whole worm lysates. Asterisks indicate non-specific bands. **(C)** Immunoblot showing expression of untagged, reencoded CYB-3 transgenes upon depletion of endogenous CYB-3 by RNAi. Actin is used as a loading control. **(D)** Viability assays for embryos depleted of *cyb-3* by RNAi and expressing the indicated reencoded *cyb-3* mutants. **(E)** Example images of embryos expressing mCherry-tagged Histone H2b and imaged under the indicated conditions. Scale bar, 5 µm. **(F)** Measurement of the NEBD to Anaphase interval in embryos imaged as in *(E)*. **(G)** Measurement of chromosome span for the indicated conditions. **(H)** *(left)* Images of representative embryos expressing WT or CDK-1 binding mutant mNeonGreen-tagged CYB-3 just prior to NEBD. Scale bar, 10 µm. *(right)* Quantification of mNeonGreen fluorescence intensity at NEBD for the indicated conditions. **(I)** Quantification of autofluorescence background-subtracted total embryo mNeonGreen::CYB-3 fluorescence after anaphase onset. Measurements were normalized to the value 20 sec prior to anaphase onset. Lanes shown in *(B)* and *(C)* are from a single exposure of the same immunoblot. *n* is the number of adult worms whose progeny were scored *(D)* or the number of embryos analyzed per condition *(F, G, H, I)*. **** represents p<0.0001 from Mann-Whitney tests.

**Figure 5:**
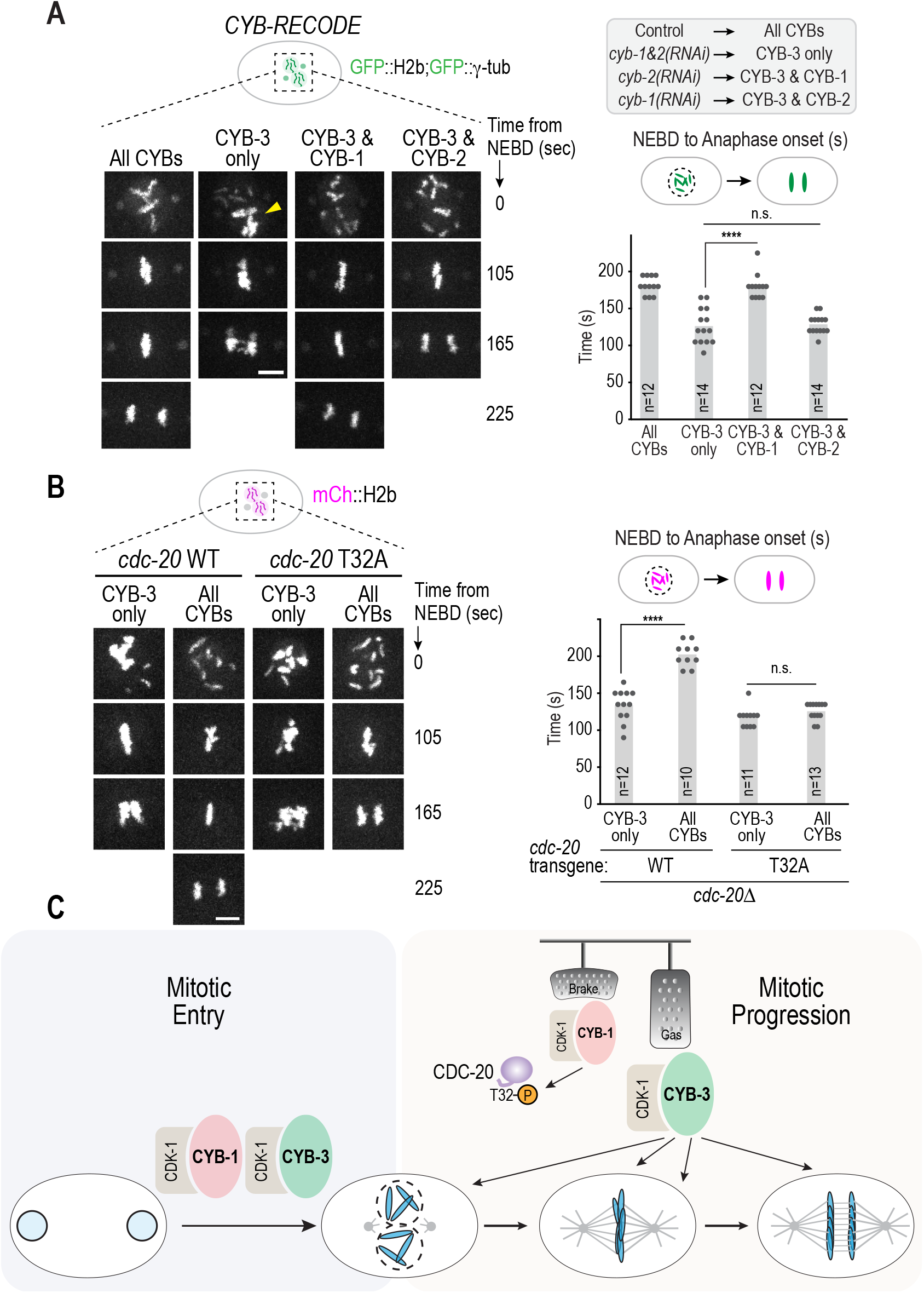
CYB-1 delays anaphase onset likely by promoting CDC-20 phosphorylation. **(A)** *(left)* Stills from timelapse sequences of *CYB-RECODE* embryos expressing GFP-tagged Histone H2b and γ-tubulin, for the conditions described in the gray box (*top right*). Yellow arrowhead points to excess chromatin caused by meiotic defects (see *Fig. S2A,B*). *(bottom right)* Quantification of the NEBD to Anaphase time interval for the specified conditions. **(B)** *(left)* Still from timelapse sequences of embryos expressing mCherry-tagged Histone H2b and either wild-type or T32A *cdc-20* in the background of a *cdc-20* null allele. *cyb-1/2(RNAi)* was used to generate the CYB-3 only condition and compare to controls having all CYBs present. *(right)* Quantification of the NEBD to Anaphase time interval for the specified conditions. Data for *cdc-20* WT and *cdc-20* T32A in the “All CYBs” condition are the same as in Fig. 3H. In all graphs, *n* represents the number of embryos analyzed. **** represents p<0.0001 from Mann-Whitney tests; non-significant (n.s.) is p>0.05. Scale bars in *(A)* and *(B)* are 5 µm. **(C)** Model summarizing key findings of the work. Both CYB-3 and CYB-1 support mitotic entry but play strikingly different roles during mitosis, schematized by the acclerator and the brake pedals, respectively. CYB-3 is the dominant cyclin B driving rapid mitosis while CYB-1 delays anaphase onset via phosphorylation of CDC-20 to ensure chromosome segregation fidelity. CYB-1 also controls spindle length (not depicted) by an as yet undefined mechanism.

We next generated a replacement system for CYB-3 in which endogenous CYB-3 is depleted by RNAi and replaced by transgene-encoded untagged CYB-3 (**Fig. S3B**). Transgene-encoded WT CYB-3 suppressed the penetrant lethality of *cyb-3(RNAi)* and rescued the significantly delayed chromosome alignment and anaphase onset in one-cell embryos (**Fig. 4C-G; S3C**). By contrast, the CYB-3 CB^Mut^ mutant recapitulated the effects of CYB-3 depletion on mitotic timing, chromosome alignment, and embryonic viability observed following CYB-3 depletion (**Fig. 4C-G**); in fact, the anaphase onset delay in CB^Mut^ CYB-3 was ∼200s longer than CYB-3 depletion (**Fig. 4F; S3C**); potentially because the mutant CYB-3 may still associate with substrates and compete with CYB-1-CDK-1 complexes that are driving the slow mitosis. These results confirm that CYB-3 associates with CDK-1 to drive rapid embryonic mitosis.

While analyzing mNG fusions WT and CB^Mut^ CYB-3, we noticed that total cellular CYB-3 levels were higher for the CB^Mut^ mutant (**Fig. 5H**). In addition, unlike WT CYB-3, CB^Mut^ CYB-3 was not degraded after anaphase onset (**Fig. 5I**). Thus, CYB-3 degradation requires association with its Cdk partner, which parallels the spindle checkpoint-independent degradation of cyclin A in early mitosis (Di Fiore and Pines, 2010; Wolthuis et al., 2008; Zhang et al., 2019).

### CYB-1 delays anaphase onset by promoting the inhibitory phosphorylation of CDC-20

In addition to showing that the pace of mitosis is slowed in CYB-1/2 only embryos, the results above indicated that the NEBD-to-anaphase onset interval in CYB-3-only embryos is ∼60s faster than in control embryos (**Fig. 2B**). The chromosome span analysis indicated that chromosome alignment occurs with similar kinetics in control and CYB-3-only embryos, but the ∼60s delay between chromosome alignment and anaphase onset observed in controls is absent in CYB-3-only embryos. To determine which cyclin(s) are responsible for this anaphase onset delay, we used the *CYB-RECODE* system to reintroduce CYB-1, CYB-2 or both (control) into CYB-3-only embryos (**Fig. 5A**). The results indicate that reintroduction of CYB-1, but not CYB-2, in CYB-3-only embryos was sufficient to restore the normal ∼60s delay between chromosome alignment and anaphase onset (**Fig. 5A; S3D**). Thus, CYB-1 introduces a delay between the completion of chromosome alignment and the onset of segregation that likely serves to ensure segregation fidelity.

Next, we next determined if CYB-1 delays anaphase onset by promoting inhibitory phosphorylation of CDC-20 on Thr32 (**Fig. 3G**). To test this possibility, we compared mitotic duration in CYB-3-only embryos expressing transgene-encoded wild-type or Thr32Ala CDC-20 in a *cdc-20*Δ background. Due to the *cdc-20*Δ allele and the recoded *cyb-1* transgene being in close genomic proximity, we were unable to do this using the *CYB-RECODE* system; however, as shown above, CYB-1 is the relevant isoform that delays anaphase onset relative to chromosome alignment in control embryos. If CYB-1 delays anaphase onset by phosphorylating CDC-20 on Thr32, then CYB-1/2 should delay anaphase onset in the presence of WT but not Thr32Ala mutant CDC-20, which is what we observed (**Fig. 5B**). We conclude that CYB-1 introduces a delay between chromosome alignment and anaphase onset by promoting the inhibitory phosphorylation of Cdc20. As shown previously, lack of this delay reduces the robustness of chromosome segregation and makes embryo viability dependent on the activity of the spindle checkpoint (Kim et al., 2017).

### Conclusion

Taken together, our results highlight a division of labor between CYB-1 and CYB-3 in controlling rapid embryonic mitoses in *C. elegans*. Cyclin B3 is the dominant fast-acting cyclin that enables rapid early embryonic cell divisions by driving mitotic processes and activation of the APC/C. At the same time, CYB-1 acts to ensure fidelity of chromosome segregation by phosphorylating CDC-20 on an inhibitory Cdk site to introduce a delay between chromosome alignment and anaphase onset. Prior work in vertebrate oocytes has highlighted a critical function for cyclin B3 in promoting the metaphase-to-anaphase transition in oocyte meiosis I by targeting the APC/C inhibitor Emi2/XErp1 for degradation (Bouftas et al., 2022; Karasu et al., 2019; Li et al., 2019). Note, however, that CYB-3 is not essential for anaphase of meiosis I in *C. elegans* oocytes (**Fig. S2A**), likely due to the absence of an Emi2/XErp1 orthologue important for metaphase II-stage oocyte arrest in vertebrates that is alleviated upon fertilization. *C. elegans* oocytes do not exhibit metaphase II arrest as their maturation and ovulation are directed by constitutive signals from sperm that are produced earlier in development and stored in the spermatheca (McCarter et al., 1999).

One model for how CYB-3 supports rapid mitosis is that the enzymatic activity of CYB-3–CDK-1 is higher than that of CYB-1–CDK1; such a model has precedence in the analysis of different cyclin-CDK complexes in yeast (Koivomagi et al., 2011; Ord et al., 2019a; Swaffer et al., 2016). A second non-exclusive model is that CYB-3 recognizes short motifs in substrates to kinetically enhance their phosphorylation, as has been demonstrated for cell cycle phase-specific cyclins in budding yeast (Bhaduri and Pryciak, 2011; Ord et al., 2020; Ord et al., 2019b). Cyclin Bs direct Cdk1 substrate specificity through two substrate recognition motifs: the hydrophobic patch and the basic patch (Loog and Morgan, 2005; Yu et al., 2021); both of these substrate-binding domains are predicted to be conserved in CYB-1 and CYB-3. With respect to CYB-1, the simplest model is that CDC-20 is a specific substrate of CYB-1–CDK-1, which would explain why adding CYB-1 to a CYB-3-only mitosis delays progression in a CDC-20 Thr32-dependent manner. We note that CYB-1–CDK-1 also controls spindle length through an unknown mechanism (**Fig. S2E**) and CYB-3–CDK-1 is needed for spindles to achieve normal microtubule density (**Fig. 3A**), highlighting additional potential favored substrates. The results presented here, which address how the central cyclin B-Cdk engine is deployed to drive rapid embryonic mitoses, offer a new opportunity to understand how distinct functional outputs are driven by different cyclin B partners of Cdk1 (**Fig. 5C**).

## MATERIALS AND METHODS

### C. elegans strains

The list of strains used in this study is in **Table S1**. *C. elegans* strains were maintained at 20°C in nematode growth media (NGM) plates containing 50 µg/mL Streptomycin and seeded with the *Escherichia coli* OP50-1 strain. For generating transgenic lines, we used the Mos1 Single-Copy Insertion (MosSCI) method (Frokjaer-Jensen et al., 2008). Briefly, transgenes were cloned into either pcFJ352 or pcFJ151 vectors and injected into strains EG6701 or EG6699 to generate single-copy insertions in chromosome I or II, respectively (Frokjaer-Jensen et al., 2008). Transformants were selected based on their ability to rescue the *unc* mobility defect of the parental strains and successful integrants were confirmed by PCR.

For the *cyb-1* and *cyb-3* RNAi replacement systems, we engineered constructs where a region of the coding sequence was modified to introduce silent mutations to render them resistant to a dsRNA targeting the endogenous gene (**Fig. S1C, S3B**). Replacement systems for CDC-20 were described previously (Kim et al., 2017).

### RNA-mediated interference (RNAi)

Double-stranded RNAs (dsRNAs) were synthesized as described before (Hattersley et al., 2018). Oligonucleotides used for dsRNA synthesis are listed in **Table S2**. dsRNAs were injected into the gut granules of L4 stage larvae at a final concentration of 0.8-1 mg/ml. Injected animals were recovered for 36-48h at 20°C before imaging. For embryonic lethality assays, L4 stage worms injected with dsRNAs were recovered at 20°C for 24h, singled onto 35 mm plates, allowed to lay progeny for an additional 24h before the mothers were removed. The next day, the total number of dead embryos and living larvae per plate were scored.

### Antibody generation

For CYB-3 antibody production, rabbits were immunized with Maltose-binding protein (MBP) fused to full-length CYB-3. Specific antibodies were affinity-purified from the serum, using a fragment containing amino acids 1-110 of CYB-3 fused to GST, to prevent the purification of antibodies against MBP (Desai et al., 2003). Antibody specificity was validated by western blot against a lysate from worms treated with a dsRNA targeting endogenous CYB-3 (**Fig. 1C**).

Because of the high amino acid sequence identity between CYB-1 and CYB-2, antibodies against these proteins were raised against their C-terminal end, using the peptides CEKINRMGQNAKVDASEME and CRMGRNLEASEAETSEME respectively. Note that the latter peptide sequence corresponds to CYB-2.2 but the antibody also detects CYB-2.1 due to their sequence similarity. Antibody specificity was confirmed by immunoblotting.

### Immunoprecipitation of mNeonGreen-tagged CYB-3

*C. elegans* strains expressing mNeonGreen-tagged CYB-3 (WT, Y-A or YIY-AAA) were seeded onto 10 mm plates and grown for 4-5 days until most of the population was adults. Animals were washed off the plates with M9 buffer (22 mM KH_2_PO_4_, 42 mM Na_2_HPO_4_, 86 mM NaCl, and 1 mM MgSO_4_•7H_2_O) and collected by centrifugation at 600g for 3 min. After two rounds of washing, pellets were resuspended in 1.5 mL of Lysis buffer (50 mM HEPES pH 7.4, 1 mM EDTA, 1 mM MgCl_2_, 100 mM KCl, 10% glycerol, 0.1% Triton X-100, 1 mM DTT, cOmplete Protease Inhibitor Cocktail (Roche)). Lysates were generated by sonication and cleared by centrifugation at 20,000g for 30 min at 4°C and incubated with mNeonGreen-nAb Agarose Beads (Allele Biotech), overnight at 4°C with rotation. Beads were washed 4 times with Lysis buffer and eluted by resuspension in 2X sample buffer (116.7 mM Tris-HCl pH 6.8, 3.3% SDS, 200 mM DTT, 10% glycerol, bromophenol blue).

### Expression and purification of proteins in Freestyle 293F cells

Binding assays were performed as described before (Ohta et al., 2021). Briefly, Myc-tagged CYB-3 (WT, Y-A or YIY-AAA) and FLAG-tagged CDK-1 constructs were expressed in FreeStyle 293-F cells (ThermoFisher Scientific). Cells were transfected using FreeStyle MAX Reagent and OptiPRO SFM according to the manufacturer’s guidelines (ThermoFisher Scientific). 20 ml of cells at a concentration of 10^6^ cells/ml were transfected with 25 µg of DNA constructs. FreeStyle 293-F cells were incubated for 48 hours at 37°C and 8% CO_2_ on an orbital shaker platform rotating at 125 rpm. 10 ml of each sample was harvested and washed with Dulbecco’s phosphate buffer saline (DPBS; ThermoFisher Scientific). Cells were then resuspended in 1 ml of lysis buffer (20 mM Tris/HCl, pH 7.5, 150 mM NaCl, 1% Triton X-100, 5 mM EGTA, 1 mM DTT, 2 mM MgCl, and one EDTA-free protease inhibitor cocktail tablet; Roche) and sonicated in an ice bath for 6 minutes. Samples were centrifuged at 13,000 rpm for 15 min at 4°C. For input samples, 50 µl of supernatant was added to 25 µl of 4x Laemmli sample buffer. The remaining 950 µl of lysate supernatant was incubated with Anti-FLAG M2 magnetic beads (Millipore Sigma) for 2 hours at 4°C. The beads were washed five times with lysis buffer and resuspended in 60 µl of 4x Laemmli sample buffer.

For immunoblotting, 4 µl of each sample was run on Mini-PROTEAN gels (Bio-Rad) and transferred to polyvinylidene difluoride (PVDF) membranes using the Trans-Blot Turbo Transfer System (Bio-Rad). Membranes were blocked with 5% nonfat dry milk in TBS-T (20 mM Tris, 150 mM NaCl, 0.1% Tween 20, pH 7.6). Membranes were rinsed with TBS-T and incubated overnight with primary antibody solutions in 5% BSA in TBS-T, followed by incubation with peroxidase-conjugated AffiniPure goat anti-mouse IgG light chain antibodies (Jackson Immunoresearch). Membranes were developed with SuperSignal West Femto substrates (ThermoFisher) and imaged using a ChemiDoc MP system (Bio-Rad). Monoclonal primary antibodies used were: anti-Flag M2 (Sigma), anti-Myc 9E10 (Sigma) and anti-α-tubulin DM1A (Millipore).

### Immunoblotting of *C. elegans* extracts

Between 40-60 adult worms per condition were transferred to tubes containing M9 buffer (22 mM KH_2_PO_4_, 42 mM Na_2_HPO_4_, 86 mM NaCl, and 1 mM MgSO_4_•7H_2_O), washed 4 times with M9 containing 0.1% Tween-20 and then resuspended in 2X sample buffer (116.7 mM Tris-HCl pH 6.8, 3.3% SDS, 200 mM DTT, 10% glycerol, bromophenol blue). After lysing by sonication followed by boiling, the equivalent of 8-10 worms was loaded onto 4-12% NuPAGE Bis-Tris Gels (Invitrogen). Proteins were then transferred to nitrocellulose membranes, probed with primary antibodies and detected using either horseradish (HRP) - conjugated secondary antibodies and WesternBright Sirius (Advansta) chemiluminescent substrate or alkaline phosphatase (AP)-conjugated secondary antibody and Western Blue® Stabilized Substrate for Alkaline Phosphatase (Promega). Membranes were imaged using a ChemiDoc MP imaging system (BioRad). Antibodies used were: rabbit anti-CYB-1, rabbit anti-CYB-2, rabbit anti-CYB-3 (all three generated in this study – *see Antibody generation*), rabbit anti-Cdk1 PSTAIRE (EMD Millipore), mouse anti-tubulin (DM1a, Sigma-Aldrich) and mouse anti-Actin (clone C4, EMD Millipore). Secondary antibodies were: HRP-conjugated donkey anti-rabbit IgG (Jackson Immunoresearch) and Alkaline Phosphatase-conjugated goat anti-mouse IgG (Promega).

### Fluorescence imaging of *C. elegans* embryos

*C. elegans* embryos were dissected from adults in M9 media and mounted onto 2% agarose pads, covered with a coverslip and imaged at 20°C. Time-lapse imaging of one-cell embryos expressing CYB-1::mNG or mNG::CYB-3 was performed on a widefield deconvolution microscope (DeltaVision Elite; Applied Precision) equipped with a charge-coupled device camera (pco.edge 5.5 sCMOS; PCO) and a 40x 1.35NA UApo/340 objective (Olympus). Single stacks were acquired at 10sec intervals with 2% illumination intensity (on an InsightSSI illuminator) and 100 ms exposure for mNeonGreen and 10% illumination intensity and 100 ms exposure for mCherry.

Time-lapse imaging of embryos and oocytes expressing GFP::tubulin and mCherry::H2b was performed on an Andor Revolution XD Confocal System (Andor Technology) with a spinning disk confocal scanner unit (CSU-10; Yokogawa) mounted on an inverted microscope (TE2000-E; Nikon), 60x 1.4 NA Plan Apochromat lenses, and outfitted with an electron multiplication back-thinned charged-coupled device camera (iXon, Andor Technology). For embryos, a 5 x 2 μm stacks were taken every 15-20 sec. Oocytes were imaged *ex utero* by cutting open adult worms on an 18 x 18 mm coverslip in 4μl of egg buffer (118 mM NaCl, 48 mM KCl, 2 mM CaCl_2_, 2 mM MgCl_2_, and 0.025 mM of HEPES, filter sterilized before HEPES addition) and directly mounting on a pad of 2% agarose in egg buffer. Oocytes were imaged in a 13 x 1 μm stack every 20 seconds. Polar body extrusion success was evaluated based on the apparent return of extruded chromosomes to the cytoplasm before the end of imaging (after meiosis II).

### Imaging analysis

For time-lapse experiments, NEBD was scored as the frame where free histone signal in the nucleus equilibrated with the cytoplasm, which was just before abrupt chromosome movements were evident. When imaging strains expressing fluorescently-labelled tubulin, NEBD was scored as the first time frame where the signal from the microtubules invaded the nuclear region. Anaphase onset was scored as the first frame with visible separation of sister chromatids.

For the measurement of chromosome span, embryo images were first oriented in the anterior-posterior axis. Signals from the Histone H2b channel were converted to binary and a bounding box was fitted to measure its width over time (**Fig. 2D**). For quantification of tubulin signal intensities, a box was drawn around the spindle area, as depicted in **Fig. 3A**. A smaller box was drawn in a region outside the spindle area to measure background fluorescence and the difference in average signal intensities between the two boxes was measured.

To quantify CYB-1::mNG and mNG::CYB-3 signal intensities over time, an area was drawn surrounding the whole embryo and the integrated density was measured; background was subtracted by copying the same area to a region without any embryos. To correct for embryo autofluorescence, we calculated baseline autofluorescence levels in a strain that expressed mCherry-tagged Histone H2b and no green fluorophore; the average autofluorescence levels were subtracted for all measurements (**Fig. S2D**).

### Statistical analyses

Statistical analysis was performed using Prism (Graphpad). In figures, asterisks denote statistical significance as calculated by Mann-Whitney tests (∗, p < 0.05; ∗∗, p < 0.01; ∗∗∗, p < 0.001; ∗∗∗∗, p < 0.0001).

## ACKNOWLEDGEMENTS

Some strains were provided by the CGC, which is funded by NIH Office of Research Infrastructure Programs (P40 OD010440). This work was supported by grants from the NIH to P.L.-G. (GM150786), A.D. (GM074215) and K.O. (GM147265). A.S. is supported by an NIH fellowship (F32GM145068). A.D. & K.O. acknowledge salary support from the Ludwig Institute for Cancer Research.

## AUTHOR CONTRIBUTIONS

P.L.-G. and A.D. initiated the project. P.L.-G., S.V., J.B. and N.V. performed the majority of the experiments. A.S. conducted imaging of meiotic embryos. A.B., S.M. and A.C.N.N. performed *in vitro* binding assays. P.L.-G., K.O. and A.D. wrote the manuscript, with input from all authors.

## Notes

### Competing Interest Statement

The authors have declared no competing interest.

