## Supplementary material for "Cyclin B3 is a dominant fast-acting cyclin that drives rapid early embryonic mitoses": Figure S1

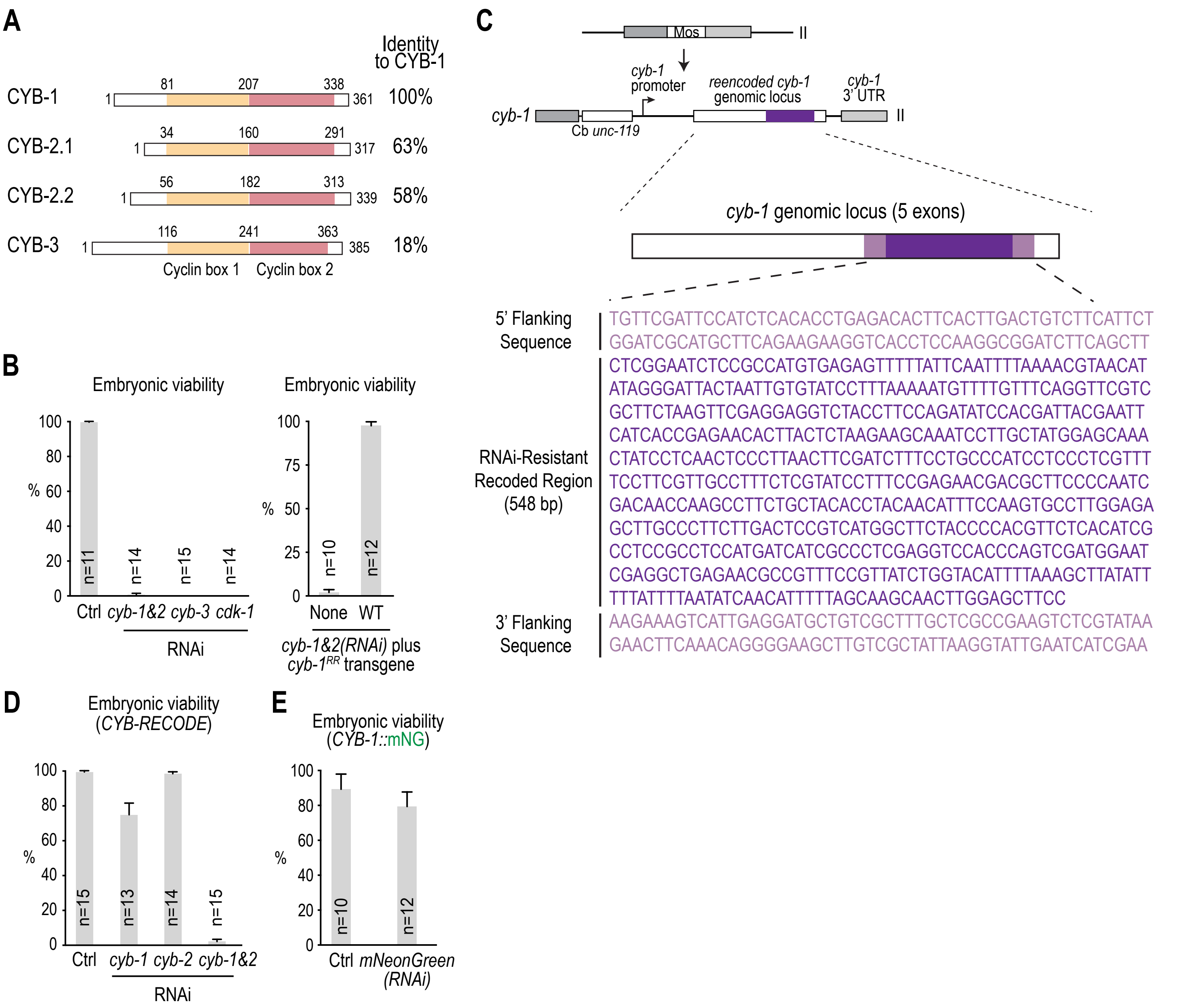


**Figure S1. Validation of the *CYB-RECODE* system.**
**(A)** Comparison of cyclin B isoforms from *C. elegans*. Numbers indicate amino acid position. **(B)** Embryonic viability measurements for the indicated conditions. **(C)** Generation of a *cyb-1* reencoded MosSCI transgene. **(D)** Embryonic viability measurements for the specified conditions. Note that expression of reencoded *cyb-1* is sufficient to fully rescue the lethality of *cyb-1&2(RNAi) (B)* but *cyb-1* and *cyb-2* are redundantly required to promote embryonic viability *(D)*. **(E)** Embryonic viability measurements for the specified RNAi conditions in a strain expressing endogenously-tagged CYB-1::mNeonGreen. Note that individual CYB-1 depletion had a mild effect on embryonic viability (panels D & E), whereas a *cyb-1*Δ mutant is lethal. The reason why the CYB-1 depletion does not exhibit lethality may be because the RNAi conditions for early embryo phenotypic analysis are optimized for depletion of the maternal load; thus, while CYB-1 is largely absent in the early embryonic divisions under these conditions, it is likely synthesized later in embryogenesis following the onset of zygotic gene expression. Error bars are 95% confidence intervals. *n* is the number of adults whose progeny was scored for viability.
