## Supplementary material for "Cyclin B3 is a dominant fast-acting cyclin that drives rapid early embryonic mitoses": Figure S2

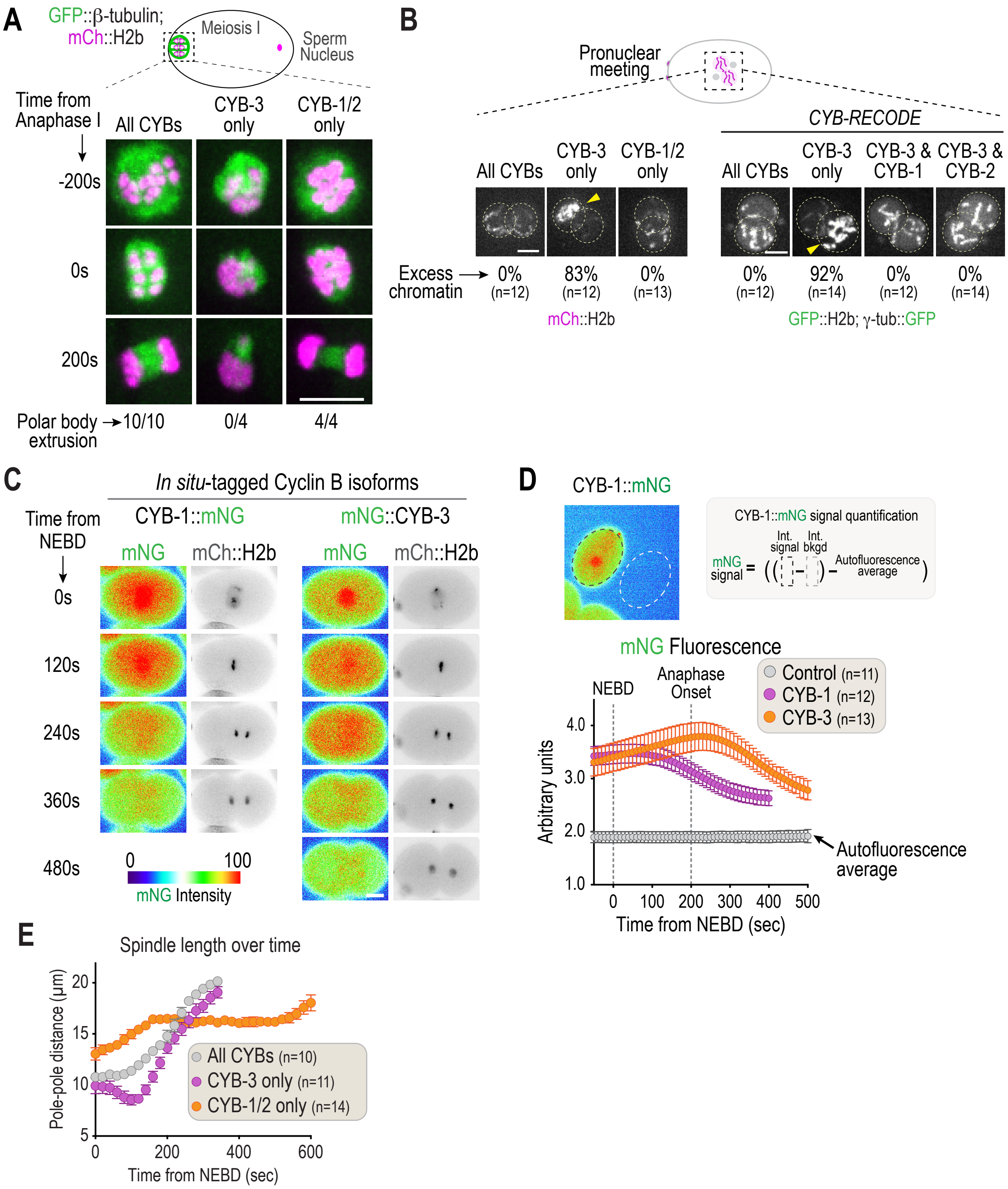


**Figure S2. Cyclin B phenotypes in meiosis and spindle elongation and analysis of cyclin B degradation kinetics in mitosis.** **(A)** Time lapse sequences of embryos undergoing meiosis I under the specified conditions. Quantification of polar body extrusion rates are included below each sequence. We note the presence of defects in spindle assembly and chromosome segregation in CYB-3-only, but not CYB-1/2-only embryos. **(B)** Example images of embryos at pronuclear meeting (75-80 seconds before NEBD), under the specified conditions. Arrow heads indicate extra chromosomal material signifying meiotic defects (*see panel A*). Numbers indicate quantification of embryos with excess chromatin for the specified conditions. **(C)** Example images of embryos expressing *in situ*-tagged cyclin B isoforms and imaged through time-lapse microscopy. Scale bar, 5 µm. **(D)** *(top)* Methodology used for the quantification of raw embryonic fluorescence intensity levels. *(bottom)* Raw fluorescence intensity levels for the specified conditions. The autofluorescence average was calculated in a strain expressing no green fluorophores and was substracted from the raw values to generate the graph in *Fig. 2F&G*. **(E)** Measurement of spindle distance over time for the indicated conditions. Whereas CYB-3-only embryos present shortening of the mitotic spindle after NEBD, CYB-1/2 embryos elongate their embryos prematurely *(see also Fig. 2A)*. Error bars are 95% confidence intervals. *n* is the number of embryos scored per condition.
