## Supplementary material for "Cyclin B3 is a dominant fast-acting cyclin that drives rapid early embryonic mitoses": Figure S3

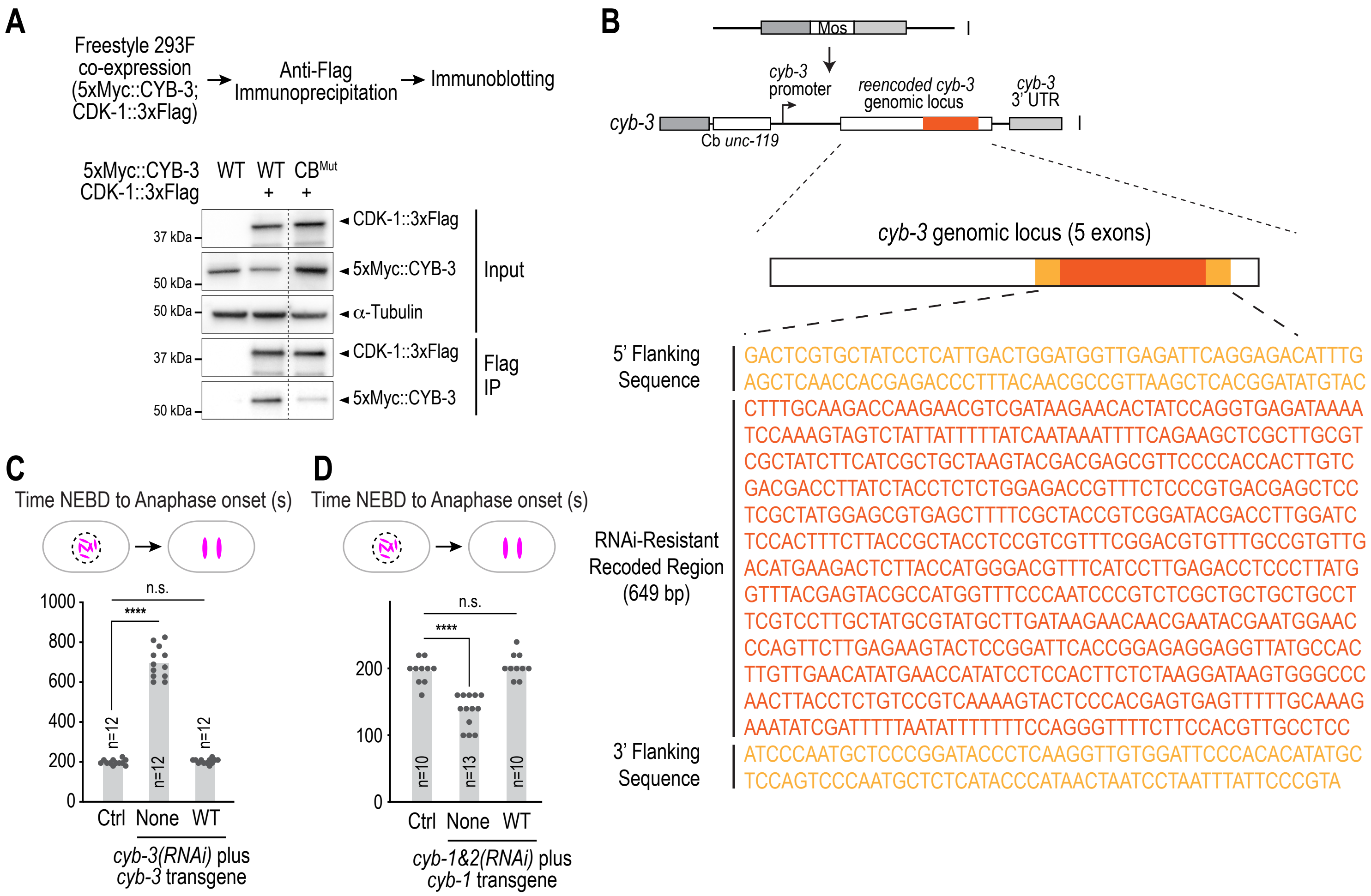


**Figure S3. Validation of CYB-3 mutants and generation of a *cyb-3* RNAi replacement system.**
**(A)** Immunoblot of an immunoprecipitation assay used to test the interaction between CDK-1 and CYB-3 mutants expressed in Freestyle 293F cells. α-Tubulin is a loading control for the input. Immunoblots were cropped from the same original image. **(B)** Strategy used to generate a *cyb-3* MosSCI transgene that was reencoded to render it resistant to *cyb-3(RNAi)*. **(C)** & **(D)** Measurement of mitotic duration in embryos imaged under the specified conditions. The *cyb-3* transgene fully rescues the mitotic duration defect of *cyb-3(RNAi) (C)*, whereas the *cyb-1* transgene is sufficient to rescue the *cyb-1&2(RNAi)* phenotype of accelerated anaphase onset *(D)*. Error bars are 95% confidence intervals. *n* is the number of embryos scored per condition. **** represents $p<0.0001$ from Mann-Whitney tests; non-significant (n.s.) is $p>0.05$.
