## Supplementary tables for "Cyclin B3 is a dominant fast-acting cyclin that drives rapid early embryonic mitoses"

**Table S1. List of strains used in this study.**

| Strain | Genotype |
|---|---|
| N2 | Ancestral |
| OD56 | *unc-119(ed3)III; ltIs37[pAA64; Ppie-1/mCherry::his-58; unc-119(+)]IV* |
| OD866 | *ltSi219[pOD1248/pSW076; Pmex-5::GFP-PH(PLC1delta1)-operon-linker-mCherry-his-11; cb-unc-119(+)]I* |
| OD868 | *ltSi220[pOD1249/pSW077; Pmex-5::GFP-tbb-2-operon-linker-mCherry-his-11; cb-unc-119(+)]I* |
| OD1702 | *unc-119(ed3)III; ltSi560 [oxTi365; oxTi365; pPLG014; Pmex-5::GFP::his-11::tbb-2_3'UTR, tbg-1::gfp::tbb-2_3'UTR; cb-unc-119(+)]V* |
| OD2729 | *ltSi1112[pOD2673/pTK049; Pcyb-1::cyb-1::mNeongreen::cyb-1 3'UTR; cb-unc-119(+)]I;unc-119(ed3)III?;ltIs37 [pAA64; pie-1/mCHERRY::his-58; unc-119 (+)] IV* |
| OD3328 | *ltSi1066[pPLG187; Pmex-5::gfp::ph::tbb-2 3'-UTR::operon linker::mCherry::his-11::tbb-2 3'-UTR; cb-unc-119(+)]II; unc-119(ed3)III* |
| OD3554 | *ltSi805[pPLG042; Pfzy-1::fzy-1::fzy-1 3'UTR; cb-unc-119(+)]I; fzy-1(lt20::loxP) ltSi1066[pPLG187; Pmex-5::gfp::ph::tbb-2 3'-UTR::operon linker::mCherry::his-11::tbb-2 3'-UTR; cb-unc-119(+)]II; unc-119(ed3)?III* |
| OD3913 | *cyb-1(lt125[cyb-1::LAP::mNeonGreen::loxP::3xFlag])IV* |
| OD4100 | *unc-119(ed3)?III; cyb-1(lt125[cyb-1::LAP::mNeonGreen::loxP::3xFlag]) ltIs37[pAA64; pie-1/mCHERRY::his-58; unc-119 (+)]IV* |
| OD4102 | *unc-119(ed3)?III; ltIs37[pAA64; pie-1/mCHERRY::his-58; unc-119 (+)]IV; cyb-3(lt135[mNeonGreen::tev::loxP::3xFlag::cyb-3])V* |
| OD4170 | *ltSi1207[pPLG270; Pmex-5::GFP-tbb-2-operon-linker-mCherry::his-11; cb-unc-119(+)]II; unc-119(ed3)III* |
| OD5083 | *ltSi1625[pPLG431; Pcyb-3::cyb-3 reencoded::cyb-3 3'UTR; cb-unc-119(+)]I; unc-119(ed3)III* |
| OD5129 | *ltSi1066[pPLG187; Pmex-5::gfp::ph::tbb-2 3'-UTR::operon linker::mCherry::his-11::tbb-2 3'-UTR; cb-unc-119(+)]II; emb-30(tn377) unc-119(ed3)?III* |
| OD5145 | *ltSi1058[pPLG149; Pfzy-1::fzy-1 T32A::fzy-1 3'UTR; cb-unc-119(+)]I; fzy-1(lt20::loxP) ltSi1066[pPLG187; Pmex-5::gfp::ph::tbb-2 3'-UTR::operon linker::mCherry::his-11::tbb-2 3'-UTR; cb-unc-119(+)]II; unc-119(ed3)III* |
| OD5149 | *ltSi1668[pPLG440; Pcyb-3::mNeonGreen::cyb-3 reencoded::cyb-3 3'UTR; cb-unc-119(+)]I; unc-119(ed3)III* |
| OD5166 | *ltSi1625[pPLG431; Pcyb-3::cyb-3 reencoded::cyb-3 3'UTR; cb-unc-119(+)]I; ltSi1066[pPLG187; Pmex-5::gfp::ph::tbb-2 3'-UTR::operon linker::mCherry::his-11::tbb-2 3'-UTR; cb-unc-119(+)]II; unc-119(ed3)III* |
| OD5191 | *ltSi1665[pPLG450; Pcyb-1::cyb-1 reencoded::cyb-1 3'UTR; cb-unc-119(+)]II; unc-119(ed3)III* |
| OD5222 | *ltSi1665[pPLG450; Pcyb-1::cyb-1 reencoded::cyb-1 3'UTR; cb-unc-119(+)]II; unc-119(ed3)?III; cyb-1(gk35)IV* |
| OD5265 | *ltSi1668[pPLG440; Pcyb-3::mNeonGreen::cyb-3 reencoded::cyb-3 3'UTR; cb-unc-119(+)]I; unc-119(ed3)III; ltIs37[pAA64; pie-1/mCHERRY::his-58; unc-119 (+)]IV* |
| OD5279 | *ltSi1712[pPLG466; Pcyb-3::cyb-3 reencoded Y112A, I116A, Y119A::cyb-3 3'UTR; cb-unc-119(+)]I; unc-119(ed3)III* |
| OD5283 | *ltSi1716[pPLG467; Pcyb-3::mNeonGreen::cyb-3 reencoded Y112A, I116A, Y119A::cyb-3 3'UTR; cb-unc-119(+)]I; unc-119(ed3)III* |

| | |
|---|---|
| OD5285 | *ltSi1712[pPLG466; Pcyb-3::cyb-3 reencoded Y112A, I116A, Y119A::cyb-3 3'UTR; cb-unc-119(+)]I; ltSi1066[pPLG187; Pmex-5::gfp::ph::tbb-2 3'-UTR::operon linker::mCherry::his-11::tbb-2 3'-UTR; cb-unc-119(+)]II; unc-119(ed3)III* |
| OD5304 | *ltSi1716[pPLG467; Pcyb-3::mNeonGreen::cyb-3 reencoded Y112A, I116A, Y119A::cyb-3 3'UTR; cb-unc-119(+)]I; unc-119(ed3)III; ltIs37[pAA64; pie-1/mCHERRY::his-58; unc-119 (+)]IV* |
| OD5326 | *ltSi1665[pPLG450; Pcyb-1::cyb-1 reencoded::cyb-1 3'UTR; cb-unc-119(+)]II; unc-119(ed3)?III; cyb-1(gk35)IV; ltSi560[oxTi365; oxTi365; pPLG014; Pmex-5::GFP::his-11::tbb-2_3'UTR, tbg-1::gfp::tbb-2_3'UTR; cb-unc-119(+)]V* |
| OD5363 | *ltSi220[pOD1249/pSW077; Pmex-5::GFP-tbb-2-operon-linker-mCherry-his-11; cb-unc-119(+)]I; ltSi1665[pPLG450; Pcyb-1::cyb-1 reencoded::cyb-1 3'UTR; cb-unc-119(+)]II; unc-119(ed3)III* |

**Table S2. Oligonucleotides used for dsRNA production.**

| Target | Oligo #1 | Oligo #2 | Template |
|---|---|---|---|
| *cyb-1&2* | taatacgactcactataggTGGAA TTTCAGCTATgtgagag | aattaaccctcactaaaggAGAGGCT CCGAGCTGTTTG | N2 genomic |
| *cyb-3* | aattaaccctcactaaaggTGTGC AAGACGAAGAATGTTG | taatacgactcactatagggCGAGGC GACATGGAAGAATA | N2 genomic |
| *cyb-1* (reencoded) | aattaaccctcactaaaggCTCG GAATCTCCGCCATGTG | TaatagactcactataggGGAAGCTC CAAGTTGCTTGC | pPLG450 |
| *cdk-1* | aattaaccctcactaaaggACCTT CCGCTCGAAACTCTT | taatacgactcactatagggTCGTAGA TCAGCAAACCctga | N2 genomic |
| *mad-3* | TAATACGACTCACTATAG Gcgaagaacttcaaaacctgga | AATTAACCCTCACTAAAGGtttgt cggtccagatccttc | N2 genomic |
| mNeonGreen | taatacgactcactataggCAAAC GACGGATACGAGGAG | aattaaccctcactaaaggCGGTGGT GTAGGACCACTTG | pPLG234 |
